# Genome degradation in the c-di-GMP signaling pathways and serovar-specific YhjH-YciG interactions rewired motility regulation in *Salmonella* Paratyphi A

**DOI:** 10.64898/2026.06.04.730116

**Authors:** Helit Cohen, Rivka Shem-Tov, Hanan Tawil, Boaz Adani, Heike Bähre, Roland Seifert, Raz Zarivach, Ohad Gal-Mor

**Affiliations:** The Infectious Diseases Research Laboratory, Sheba Medical Center, Tel-Hashomer, Israel; Faculty of Medical &Health Sciences, Tel Aviv University, Tel Aviv, Israel; Department of Clinical Microbiology and Immunology, Tel Aviv University, Tel Aviv, Israel; Research Core Unit Metabolomics, Hannover Medical School, Hannover, Germany; Institute of Pharmacology & RCU Metabolomics, Hannover Medical School, Hannover, Germany; Departments of Life Sciences, National Institute for Biotechnology in the Negev (NIBN), Ben-Gurion University of the Negev, Be’er Sheva, Israel

**Author notes:** These authors contributed equally to this work (Helit Cohen, Rivka Shem-Tov). Author for Correspondence: Ohad Gal-Mor, The Infectious Diseases Research Laboratory, Sheba Medical Center, Tel-Hashomer, Israel.

**Keywords:** *Salmonella*, c-di-GMP, YhjH, YcgR, YciG, motility, flagella, Paratyphi A, genome degradation

## Abstract

*Salmonella enterica* serovars Typhimurium (STM) and Paratyphi A (SPA) cause clinically distinct diseases, yet the molecular bases for their different lifestyles remain incompletely understood. Genome degradation, a hallmark of typhoidal *Salmonella*, results in extensive pseudogenization across multiple functional pathways. Here, we investigate how such gene inactivation rewires cyclic-di-GMP (c-di-GMP) signaling and flagellar motility regulation in SPA vs. STM. We show that YhjH, a conserved phosphodiesterase (PDE), is required for motility in STM but not in SPA, despite retaining PDE activity in both serovars. We demonstrate that this functional divergence is caused by pseudogenization of *ycgR* in SPA, which truncates the flagellar brake protein YcgR to a nonfunctional peptide, severing the link between c-di-GMP levels and flagellar motor control. Site directed mutagenesis in the YhjH active site and ectopic expression of intact YcgR from STM that restored YhjH-dependent motility regulation in SPA, confirmed this molecular mechanism. Additionally, using a bacterial two-hybrid (BACTH) genetic screen, we identified a serovar-specific interaction between YhjH and the general stress protein YciG in SPA, but not in STM. Computational RNA folding analysis predicted substantial differences in mRNA secondary structure and stability between the SPA and STM *yhjH* alleles, suggesting a potential role for synonymous mutations in this serovar-specific interaction. Together, these findings reveal how genome degradation can rewire regulatory networks, uncovering a fundamental difference in motility control between typhoidal and non-typhoidal *Salmonella* and suggest that these differences allow SPA motility under conditions that suppress motility in STM.

**IMPORTANCE:** *Salmonella enterica* serovars Typhimurium (STM) and Paratyphi A (SPA) cause fundamentally different diseases in humans, yet the molecular basis for their distinct lifestyles and pathogenicity remains poorly understood. Genome degradation is a hallmark of typhoidal *Salmonella*, but its functional consequences for regulatory networks are largely unexplored. Here, we demonstrate that pseudogenization of *ycgR* in SPA dismantles c-di-GMP-mediated flagellar motor control, liberating SPA from an inhibitory brake that suppresses motility in STM. Additionally, we uncover a serovar-specific interaction between the phosphodiesterase YhjH and the general stress protein YciG in SPA, demonstrating that genome degradation can drive regulatory network rewiring beyond simple gene loss. These differences in motility regulation may facilitate the systemic pathogenesis and unique lifestyle of SPA.

## INTRODUCTION

*Salmonella enterica* is a ubiquitous and diverse Gram-negative food-borne and zoonotic bacterial pathogen. This single species consists of more than 2600 antigenically-distinct serovars (1) that can colonize the gut and cause different diseases in a wide range of hosts, including wild and food-producing animals as well as humans (2, 3). Different *S. enterica* serovars that infect humans can be functionally classified into two groups according to the clinical manifestations and the disease they cause. Typhoidal *Salmonella* serovars that include *S. enterica* serovar Typhi (*S*. Typhi), *S. enterica* serovar Paratyphi A (*S*. Paratyphi A or SPA from hereinafter) and *S. enterica* serovar Sendai (*S*. Sendai) are all human-specific serovars that can cause a life-threatening febrile disease known as enteric (or typhoid) fever (4). This is a non-inflammatory systemic disease, in which the pathogen reaches the blood circulatory system and colonizes the mesenteric lymph nodes, liver, spleen, and sometimes even the nervous system (5).

In contrast, dozens of non-typhoidal *Salmonella* (NTS) serovars including *S. enterica* serovar Typhimurium (*S*. Typhimurium or STM from hereinafter) are generalist pathogens with a wide range host-specificity that can infect different reptile, amphibian, avian and mammalian hosts and cause an acute gastroenteritis disease in immunocompetent humans (6). In most cases, this is a self-limited inflammatory disease of the terminal ileum and colon, which does not proceed to the dissemination of the pathogen behind the intestinal submucosa (7). *S. enterica* infections still pose a significant health challenge worldwide, with an annual of 27 million cases of enteric fever (8) and 78.7 million infections of NTS-mediated gastroenteritis (9), globally.

Although it is well-known that typhoidal and NTS serovars cause clinically different diseases (7), the molecular and cellular mechanisms responsible for their distinct disease manifestations remained elusive. This gap in knowledge is especially relevant for SPA that does not express the Vi capsule as *S*. Typhi, which plays a key role in pathogen modulation of the host immune response (10). Nonetheless, a common genetic signature, which is shared by all typhoidal serovars, is the phenomenon of genome degradation. This process of genomic decay involves multiple events of gene inactivation (pseudogene formation) as well as complete loss or gene deletion from the genome (11–14). As a result, while STM harbors very low portion of pseudogenes in its genome (15), nearly 5% of the annotated coding sequences (CDS) in *S*. Typhi and SPA are pseudogenes (16–18). Therefore, genome degradation is thought to affect and play, in a yet not fully understood manner, a role in the distinct lifestyle of typhoidal serovars (17, 19).

Cyclic diguanylate monophosphate (c-di-GMP) is an abundant bacterial nucleotide-based second messenger (20). In recent years, c-di-GMP has emerged as a key regulator of bacterial physiology and virulence that controls multiple cellular processes, including the switch between motile and nonmotile state and the transition between acute and persistent infection (21, 22). Cellular c-di-GMP concentration is determined by the function of c-di-GMP turnover proteins including diguanylate cyclases (DGCs) that synthesize c-di-GMP from GTP through their catalytic GGDEF domains, and c-di-GMP-specific phosphodiesterases (PDEs), which degrade c-di-GMP and harbor either the EAL or the HD-GYP domain (23). One of the main phenotypes derived by high c-di-GMP cellular concentration is the inhibition of flagellar-based motility (24). This occurs by c-di-GMP binding to the PilZ domain of the flagellar brake effector protein YcgR, which results in a conformational change and the direct binding of the YcgR-c-di-GMP complex to the flagellar motor (25).

STM encodes as many as 22 GGDEF/EAL domain proteins with a putative c-di-GMP metabolizing activity (26). Nevertheless, YhjH (AKA STM3611 or PdeH) is considered to be the most active PDE that was shown to play a role in *Salmonella* motility regulation (27, 28). YhjH is also a member of the flagella-chemotaxis regulon and is expressed from a flagellar class III promoter, which is regulated by the alternative sigma factor FliA (σ^28^) (29). Moreover, it was reported that YhjH overexpression has led to elevation in both modes of motility, swimming (unicellular motion) and swarming (multicellular motion) (30), while a deletion of *yhjH* resulted in a significantly suppressed motility (29, 31).

In this study, we show that unexpectedly, YhjH plays a distinct role in motility regulation in STM vs. SPA. We demonstrate that YhjH fails to regulate motility in SPA due to an *ycgR* gene inactivation that is truncated in SPA, but intact in STM. Moreover, we found that in SPA, but not in STM, YhjH interacts with YciG, a conserved general stress protein, with a previously unknown function. Overall, our findings demonstrate how genome degradation in the c-di-GMP regulatory network rewires flagella-based motility regulation in the clinically important pathogen, SPA and suggest that such changes may contribute to its unique lifestyle and pathogenicity.

## RESULTS

### YhjH regulates motility in STM but not in SPA

YhjH is a highly conserved 255 amino acid protein in STM and in SPA, with 99.2% identity on the protein level (Fig. 1A). Previous studies have reported a central role for YhjH in motility regulation in STM (29, 31), however, to the best of our knowledge, the contribution of YhjH to motility regulation in SPA has never been reported. Considering differences in the global regulation of the flagella-motility regulon between STM and SPA, as we have previously reported (32–34), we sought to compare the role of YhjH in motility in both serovars. To address that, an identical in-frame non-polar deletion was constructed in the *yhjH* gene in SPA (from hereafter we will refer to this variant as *yhjH*_SPA_) and STM (from hereafter we will refer to this variant as *yhjH*_STM_), and the swimming motility of both null strains was studied on semisolid agar plates at 37°C. Interestingly, while the deletion of *yhjH* (Δ*yhjH*) in STM significantly reduced its motility, the homologous deletion in SPA, did not affect its motility, which is generally much lower in SPA than in STM (Fig. 1B and C). Complementing Δ*yhjH* in STM, by expressing the full *yhjH* gene either as *yhjH*_STM_ or *yhjH*_SPA_ variant from a low copy number vector (pWSK29), restored the motility level of the STM Δ*yhjH* strain to similar motility levels as in the WT parental strain (Fig. 1D and E). We concluded from these experiments that YhjH in STM, but not in SPA, regulates flagella-mediated motility and that, because YhjH_SPA_ can fully replace the YhjH_STM_ allele, these differences are not due to amino acid variations (at positions 192 and 199), found in the YhjH sequence between STM and SPA (Fig. 1A).

**Figure 1.**
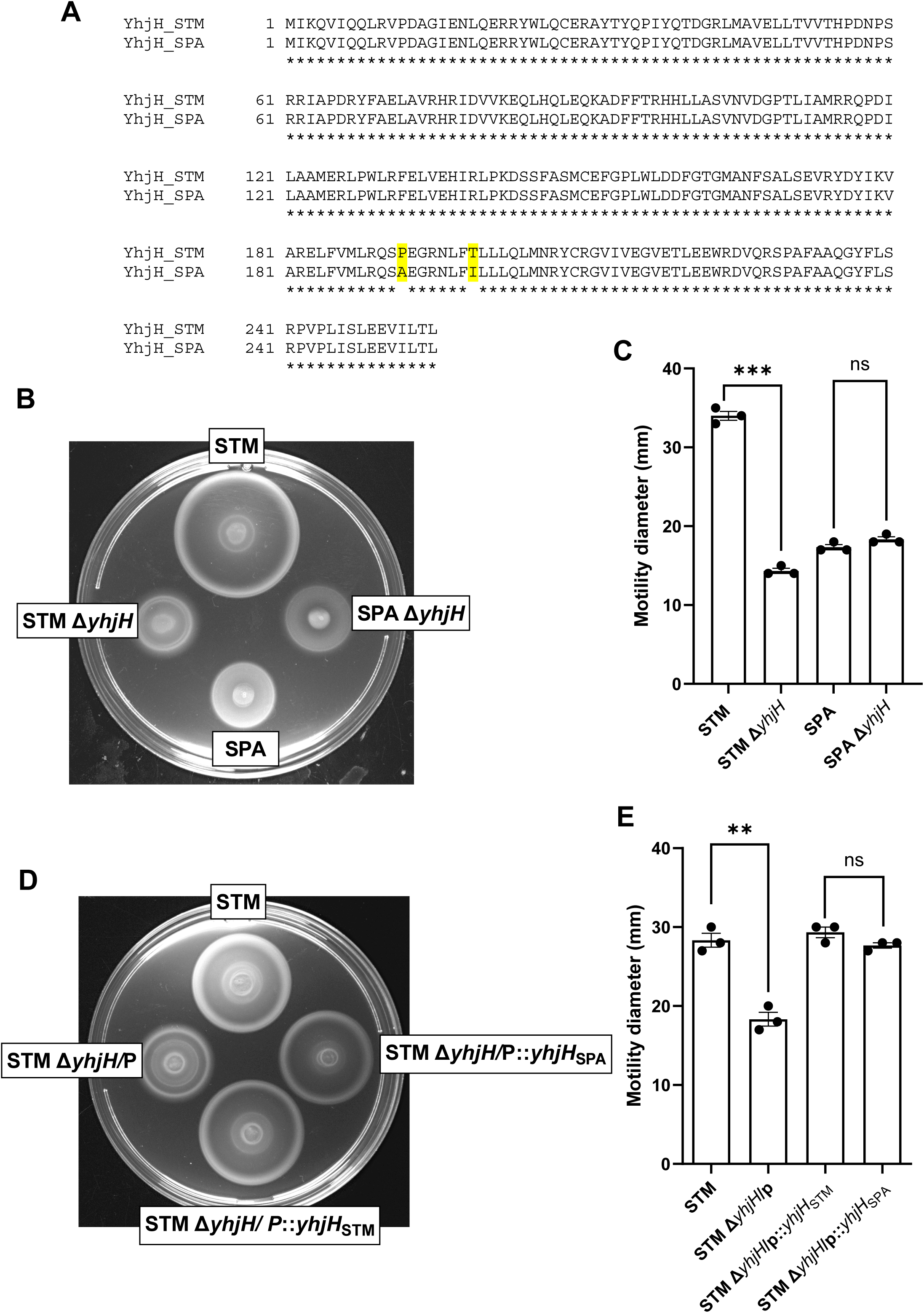
YhjH plays a role in motility regulation in STM but not in SPA. (**A**) Pairwise comparison of the amino acid sequence of the YhjH orthologs in STM (accession number CBW19670) and SPA (accession number AAV79276 identical in strains 45157 and ATCC 9150). Divergent amino acids at positions 192 and 199 are highlight. (**B**) Cultures of WT STM and SPA and their isogenic Δ*yhjH* strains were grown at 37 °C aerobically for overnight and spotted on soft LB agar plates that were incubated at 37 °C for 3.5 h. **(C)** The measured swimming radius of the cultures at 37 °C is shown. Bars indicate the mean value and its associated standard error of the mean (SEM) of three independent biological repeats. (**D**) STM wildtype strain and its isogenic Δ*yhjH* null strain that was complemented with a pWSK29 empty vector (STM Δ*yhjH*/P), pWSK29 expressing *yhjH* from STM (STM Δ*yhjH*/P::*yhjH*_STM_) or pWSK29 expressing *yhjH* from SPA (STM Δ*yhjH*/P::*yhjH*_SPA_) were grown and assayed as above. (**E**) The measured swimming radius of the cultures at 37 °C is shown. Bars indicate the mean value and its associated SEM of three independent biological repeats. An Unpaired, 2-tailed Student *t* test was used to calculate statistical significance between compared strains. **, P-value <0.01; ***, P-value < 0.001; ns, not significant.

### YhjH suppresses biofilm formation in SPA

In addition to its role in motility control, YhjH was previously demonstrated to act as a negative regulator of biofilm in STM (31, 35), and ectopic expression of STM_YhjH_ in closely related species was shown to reduce biofilm production (36, 37). Since we found that in SPA, YhjH does not regulate motility, we next asked whether YhjH still plays a role in biofilm formation in this serovar. To this end we grow SPA and STM cultures under biofilm inducing conditions (at 28°C in LB medium depleted of NaCl) and determined the biofilm mass generated in the WT background compared to its isogenic Δ*yhjH* strain. Interestingly, while biofilm in general was found to be produced at much lower levels in SPA than in STM, the lack of *yhjH* significantly enhanced biofilm production in SPA cultures grown under biofilm inducing conditions. Complementation of *yhjH* from a low-copy-number plasmid (pWSK29::*yhjH*_SPA_) reduced biofilm formation to similar levels as in the SPA wildtype background (Fig. 2). We concluded from this experiment that while YhjH does not regulate motility in SPA, it is still produced as a functional protein that plays a role as a negative regulator of biofilm formation in this serovar.

**Figure 2.**
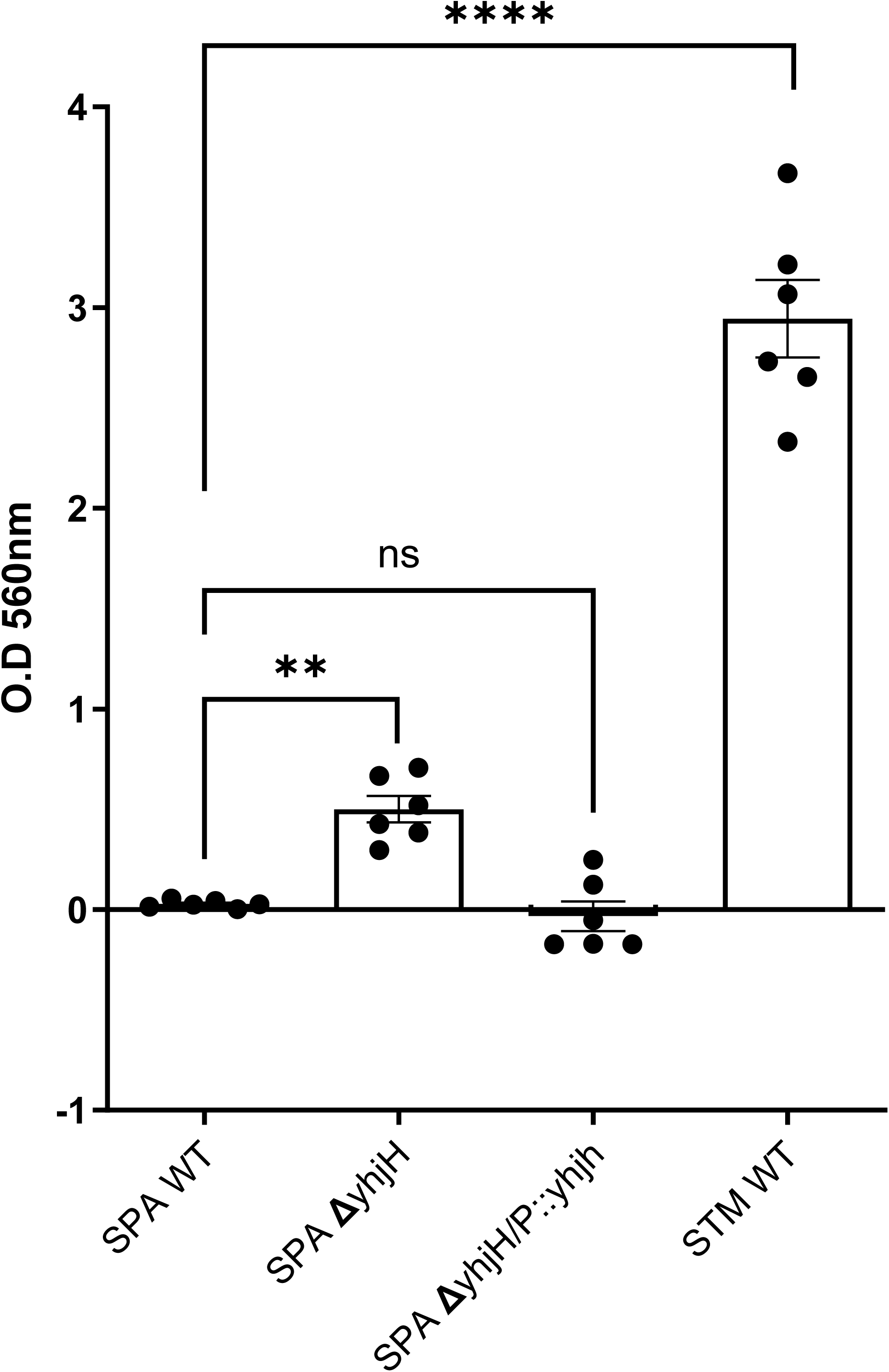
YhjH suppresses biofilm formation in SPA. Biofilm formation by WT SPA, its isogenic Δ*yhjH* strain, Δ*yhjH* complemented with pWSK29::*yhjH* and WT STM was determined by Crystal Violet staining of static cultures that were incubated for 96 h at 28 °C in rich LB broth in the absence of sodium chloride (biofilm induced conditions). Biofilm was quantified by the absorbance of the stained biomass at OD_560_. The bars show the mean of six biological repeats and the SEM is indicated by the error bars. One-Way ANOVA with Dunnett’s multiple comparisons test was used to determined statistical significance relative to SPA WT. **, P-value <0.01; ****, P-value < 0.0001; ns, not significant.

### YhjH functions as a PDE in both STM and SPA

The distinct role of YhjH in flagella-based motility in STM, but not in SPA, has led us to examine its potential activity as phosphodiesterase in both serovars. To do that we first determined the c-di-GMP concentration in STM and SPA and in their Δ*yhjH* isogenic strains. We reasoned that if YhjH functions as a dominant PDE, the cellular c-di-GMP concentration is expected to be higher in the Δ*yhjH* background, due to lower degradation of c-di-GMP in these strains. Indeed, quantification of cytosolic c-di-GMP concentration by mass spectrometry in samples extracted from STM and SPA cultures that were grown in LB medium to the stationary phase at 30°C, indicated that the absence of *yhjH* in both serovars was associated with a significant increase in c-di-GMP concentration in both serovars (Fig. 3A). These results suggested that YhjH functions as a key PDE both in SPA and STM.

**Figure 3.**
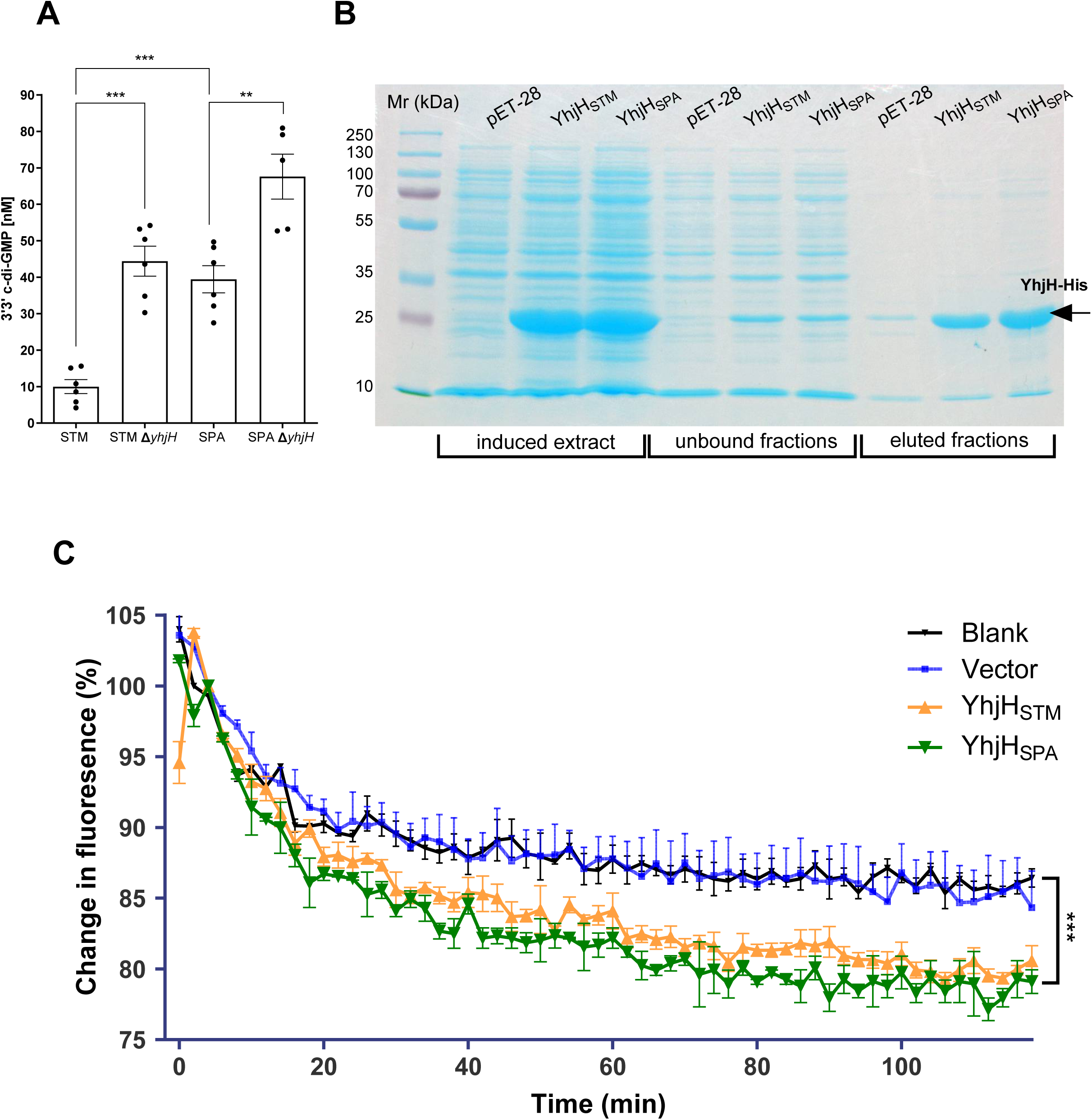
YhjH has a phosphodiesterase activity in both STM and SPA. (**A**) Wildtype STM and SPA and their isogenic Δ*yhjH* mutant strains were grown in liquid cultures overnight at 30 °C and subjected to cyclic dinucleotides extraction. The concentration of c-di-GMP in the extracted fractions was analytically measured using high-performance liquid chromatography-tandem mass spectrometry. The mean value of c-di-GMP concentration and its SEM are shown for 5-6 independent cultures. An Unpaired, 2-tailed Student *t* test was used to calculate statistical significance between compared strains. **, P-value <0.01; ***, P-value < 0.001. (**B**) The coding sequence of *yhjH* from STM (YhjH_STM_) and SPA (YhjH_SPA_) were cloned and expressed in pET-28a vector, to generate a recombinant YhjH protein with a six-histidine sequence (6xHis) added to its carboxyl-terminus under IPTG inducing promoter. The recombinant tagged proteins were induced and purified on nickel beads. Fractions (10 µL) of the purified proteins and the different stages of the expression and furification are shown after separation on a 12% SDS-PAGE and Coomassie Brilliant Blue staining. (**C**) The phosphodiesterase activity of the purified His-tagged YhjH_STM_ and YhjH_SPA_ proteins (42 pmol/ 1.2 µg) was measured by monitoring the change in the fluorescence of MANT-labeled cyclic di-GMP substrate at 37 °C over time. The substrate MANT-c-di-GMP only (without protein; Blank) and substrate that was incubated with equivalent volume (8 µL) of eluted extract from culture expressing the empty pET-28a plasmid (Vector) were included as negative controls representing spontaneous degradation of the substrate. Each time point shows the mean and SEM of three wells. An Unpaired, 2-tailed Student *t* test was used to calculate statistical significance between reactions. ***, P-value < 0.001.

To further confirm the PDE activity of YhjH in STM and SPA, we expressed and purified recombinant His-tagged YhjH proteins from STM and SPA. To this end, the intact *yhjH* gene from both serovars was PCR amplified and cloned into the pET-28a vector, generating a C-terminal hexahistidine-tagged version of YhjH_STM_ and YhjH_SPA_. The quality and the purity of the tagged proteins following purification on Ni-NTA magnetic beads were analyzed by sodium dodecyl sulfate polyacrylamide gel electrophoresis (SDS-PAGE) that showed strong induction and enriched purification of recombinant YhjH tagged proteins in the eluted fractions (Fig. 3B). Subsequently, their PDE activity was estimated in vitro by monitoring the change in a fluorescence cyclic di-GMP subtract that was MANT-labeled (Di-MANT-c-di-GMP). These assays demonstrated a similar rate of decrease in the labeled c-di-GMP substrate in the presence of 42 pmole of either YhjH_STM_-His or YhjH_SPA_-His, which was significantly catalyzed compared to a spontaneous degradation of this substrate (Fig. 3C). Collectively, we concluded from these experiments that although YhjH plays a different role in motility regulation in STM and SPA, in both serovars, YhjH functions as a major PDE that can catalyze c-di-GMP breakdown both *in vitro* and in *Salmonella* cells.

### YhjH motility regulation is dependent on its PDE activity

Next, we asked whether the PDE activity of YhjH is required for its motility regulation in STM. Previous work has shown that a glutamate to alanine substitution in position 136 (E136A) at the active site of YhjH leads to a loss of its PDE activity (31, 36). Therefore, we used site-directed mutagenesis and constructed a catalytically inactive derivative of YhjH in the STM background and tested its effect on swimming motility. As shown in Fig. 4, the E136A substitution (YhjH_E136A_) abolished the swimming ability of STM to a level similar to a null Δ*yhjH* mutant. The impaired motility could be complemented by an ectopic expression of the *yhjH*_STM_ allele in the *yhjH*_E136A_ mutant strain (Fig. 4A-B).

**Figure 4.**
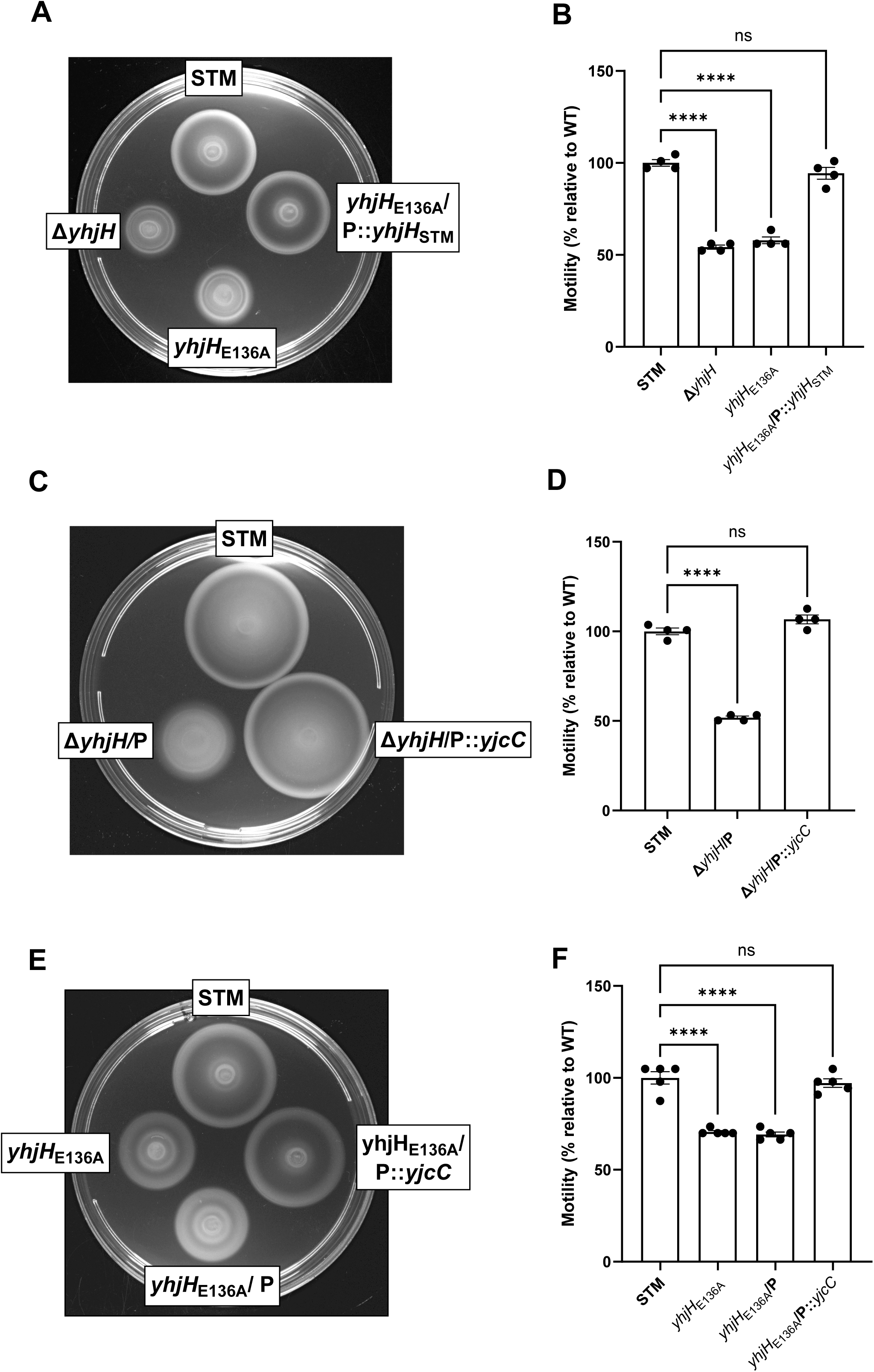
YhjH-mediated motility regulation is dependent on its PDE activity. **(A)** An STM WT strain, its isogenic Δ*yhjH* strain, a mutant harboring an E136A YhjH substitution (*yhjH*_E136A_) and a complementing strain harboring the WT STM YhjH copy on a plasmid (*yhjH*_E136A_/P::*yhjH*_STM_) were subcultured at 37 °C in LB and aliquots of 10 µL were placed onto soft agar plate. The swimming motility of these strains was measured after incubation for 3 h at 37 °C. (**B**) The measured swimming radius of the cultures relative to the motility of the WT background (that was normalized to 100%) at 37 °C is shown. Bars indicate the mean value and its associated SEM of four independent biological repeats. (**C**) An STM wildtype strain, its isogenic Δ*yhjH* l mutant strain, harboring an empty pBAD18 plasmid (Δ*yhjH*/P) or complemented with pBAD expressing the PDE encoded gene *yjcC* (ΔyhjH/P::*yjcC*) cultures were subcultured in LB supplemented with 10 mM arabinose at 37 °C and placed onto soft agar plates that were supplemented with 10 µg/ mL arabinose, which were incubated for 3 h. (**D**) The measured swimming radius of the cultures relative to the motility of the WT background (that was normalized to 100%) at 37 °C is shown. Bars indicate the mean value and SEM of four independent biological repeats. (**E**) An STM WT strain and its isogenic *yhjH* mutant carrying an E136A substitution *(yhjH*_E136A_) and this mutant strain complemented with an empty pBAD18 vector or with pBAD::*yjcC* were grown and assayed as above. (**F**) The measured swimming radius of the cultures relative to the motility of the WT background (that was normalized to 100%) at 37 °C is shown. Bars indicate the mean value and its associated SEM of five independent biological repeats. Statistical significance was determined by one-way ANOVA followed by *post hoc* Dunnett’s test. ****, P-value < 0.0001; ns, not significant.

Moreover, overexpression of a different *Salmonella* PDE coding gene, *yjcC* (38) that was expressed under an arabinose-inducible promoter was able to complement both Δ*yhjH* deletion (Fig. 4C-D) and the *yhjH*_E136A_ mutation (Fig. 4E-F), and restored their motility to a similar level as in the WT STM background. Accumulatively, we concluded from these experiments that motility regulation in STM by YhjH is mediated by its PDE activity and that a YhjH loss of function can be compensated by an overexpression of an alternative PDE.

### YhjH fails to control SPA motility due to a YcgR inactivation

Since we found that YhjH in SPA retains its PDE activity, we hypothesized that its inability to regulate SPA motility lies further downstream along its regulatory cascade. In STM, it was shown that the c-di-GMP-binding protein YcgR acts as a "molecular brake" that interacts with the motor proteins MotA and FliG to reduce motor speed in response to high c-di-GMP levels (39, 40). Upon c-di-GMP binding, YcgR decreases motor speed and biases rotation towards counterclockwise, thus promoting the transition from a motile to sessile state. Therefore, the lack of the PDE, YhjH results in elevated cellular concentrations of c-di-GMP and a slower flagellar spinning that reduces motility (24, 25). Interestingly, when we analyzed the *ycgR* loci in the SPA genome, we found that, in contrast to STM, *ycgR* is a pseudogene in SPA due to a frameshift mutation, leading to a truncated peptide of only 13 amino acids (Fig. 5A).

**Figure 5.**
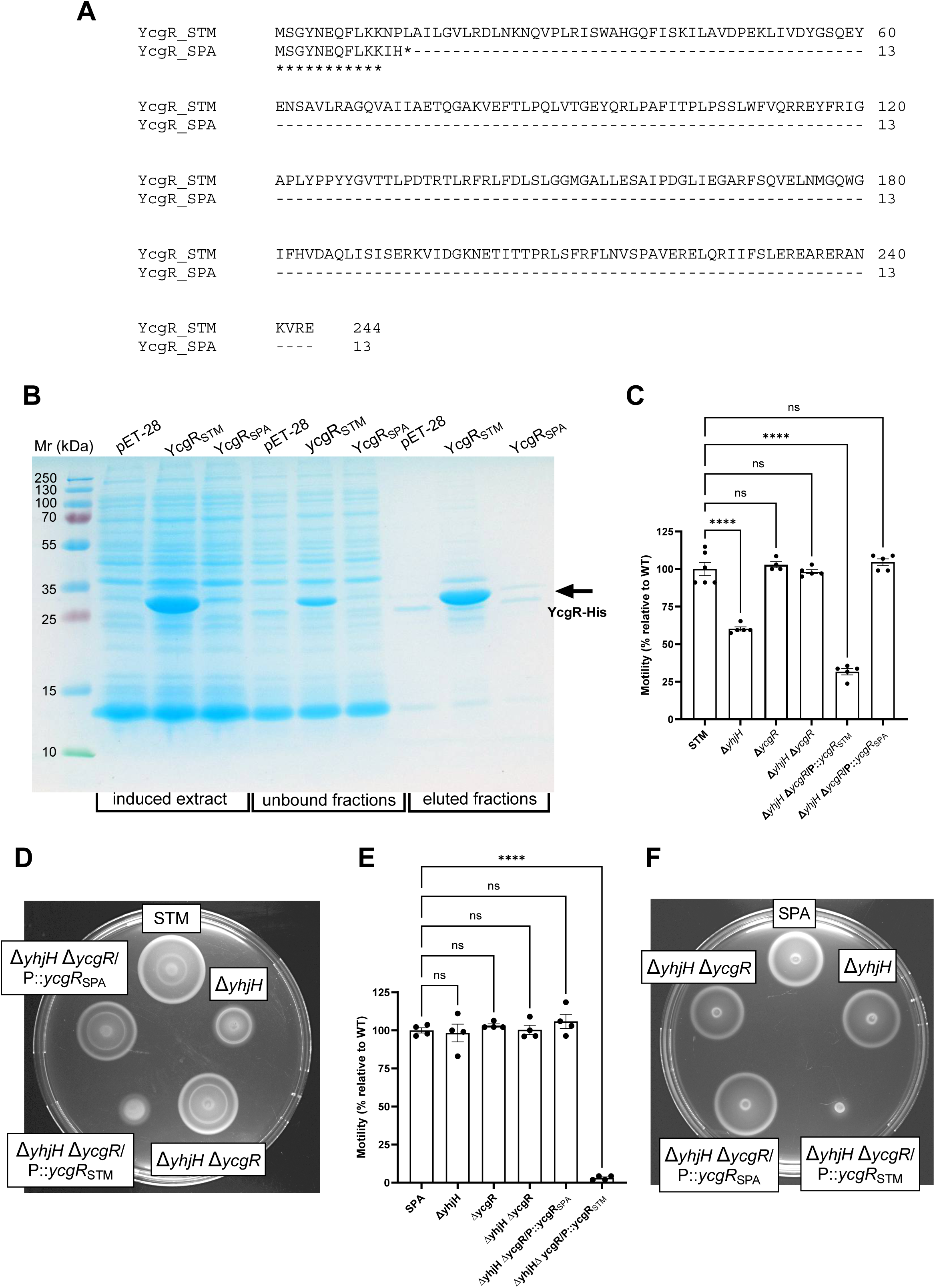
YhjH fails to control SPA motility due to YcgR inactivation. (**A**) The *ycgR* orthologous genes from STM (accession number CBW19670) and SPA (accession number AAV79276, identical in SPA 45157 and ATCC 9150) were translated and aligned using the Clustal Omega alignment tool. A termination at the 14^th^ codon in the SPA YcgR is indicated by a highlighted asterisk. (**B**) The ycgR coding sequences from STM and SPA were cloned in pET-28a to generate a recombinant YcgR protein with a six-histidine sequence (6xHis) added to its N-terminus, under IPTG inducing promoter. The recombinant tagged proteins were induced and purified on nickel beads. Fractions (10 µL) of the purified proteins and aliquots obtained during the different stages of the expression and extraction are shown after separation on a 15% SDS-PAGE and Coomassie Brilliant Blue staining. (**C**) An STM wildtype strain, its isogenic single mutant strains Δ*yhjH, ΔycgR,* a double Δ*yhjH ΔycgR* mutant strain and a double Δ*yhjH ΔycgR* mutant strain complemented with pWSK29 plasmid carrying the *ycgR* gene from SPA (P::*ycgR*_SPA_) or STM (P::*ycgR*_STM_) were grown overnight at 37 °C in liquid cultures and placed onto soft agar plate. The swimming motility of these strains was measured after incubation for 3 h at 37°C. Bars indicate the mean value and SEM of 4-6 independent biological repeats. (**D**) A representing image of this motility assay with relevant STM strains is shown. (**E**) A SPA WT strain, its isogenic mutant strains Δ*yhjH*, Δ*ycgR*, a double Δ*yhjH ΔycgR* mutant strain and a double Δ*yhjH ΔycgR* mutant strain complemented with pWSK29 plasmid carrying the *ycgR* gene from SPA (P::*ycgR*_SPA_) or STM (P::*ycgR*_STM_) were grown and assayed as above. The swimming motility of these strains was measured after incubation for 3 h at 37 °C. Bars indicate the mean value and SEM of four independent biological repeats. (**F**) A representing image of this motility assay with relevant SPA strains is shown.

To exclude the possibility that an altered version of YcgR is still being translated in SPA, via an alternative start codon, we cloned the native *ycgR* open reading frame from STM (*ycgR*_STM_) and SPA (*ycgR*_SPA_) into the expression vector pET-28a, while generating a 6×His-tag at its N-terminus and expressed these constructs in *Escherichia coli* BL-21 cells. This surrogate expression indicated a robust expression of His-YcgR_STM_, but the lack of any detectable expression of a tagged His-YcgR_SPA_ (Fig. 5B). We concluded from these experiments that while YcgR is intact and expressed in STM, its homolog in SPA is inactive and not translated into a mature protein.

To test whether the inability of YhjH to regulate motility in SPA is due to an inactive YcgR, a single null in-frame Δ*ycgR* deletion and a double Δ*yhjH* Δ*ycgR* deletion were constructed in the STM and SPA backgrounds. A Δ*ycgR* or a double Δ*yhjH ΔycgR* deletion did not have a significant effect on STM (Fig. 5C and D) or SPA (Fig. 5E and F) motility. Nevertheless, when the Δ*yhjH ΔycgR* mutants in STM or in SPA were complemented by ectopic expression of YcgR_STM_, the motility of both strains was dramatically reduced. In contrast, no change in the motility was identified when the Δ*yhjH ΔycgR* mutants were complemented with the YcgR_SPA_ (Fig. 5C-F). We therefore concluded from these experiments that the *ycgR* gene in SPA is degraded and translated into an inactive protein that prevents YhjH motility regulation in SPA. Replacing the inactivated YcgR protein in SPA with an active ortholog from STM restored the ability of the SPA YhjH to regulate motility in SPA in a similar manner as in STM.

### YhjH interacts with the general stress protein YciG in SPA, but not in STM

To further study potential differences in YhjH activity in SPA and STM, we next sought to identify YhjH-interacting proteins using a bacterial two-hybrid (BACTH) approach. This system is based on adenylate cyclase (AC) reconstitution in an *E. coli* strain, lacking endogenous adenylate cyclase activity and exploits the fact that the catalytic domain of *Bordetella pertussis* AC (CyaA) is composed of two complementary domains (T25 and T18) that are inactive when they are physically separated (41, 42). If T18 and T25 are fused to two interacting proteins, their interaction will cause a functional complementation by bringing to proximity the T25 and T18 domains. As a result, cyclic AMP produced by the reconstituted chimeric AC enzyme binds to the catabolite activator protein CAP, which in turn induces the expression of the *lac* operon and the catalytic activity of β-galactosidase. The β-galactosidase enzymatic activity is therefore correlated with cAMP levels produced in the cells and can be visually detected by the production of a blue pigment in the presence of the X-gal (5-bromo-4-chloro-3-indolyl-beta-D-galacto-pyranoside) substrate. Consequently, the production of blue pigment by *E. coli* colonies indicates interaction between the polypeptides fused to T25 and T18, and the activity level of β-galactosidase is proportionate to the degree of protein-protein interaction between the T-18 and T-25 fused proteins (42, 43).

Building on this approach, we used the BACTH system to identify SPA proteins potentially interacting with YhjH (Fig. 6A). To this end, we fused the YhjH_SPA_ to the T25 domain and transformed this construct (pKT25::*yhjH*_SPA_) into an *E. coli cya* mutant strain that requires plasmid-derived AC activity for the activation of the *lac* operon (*E. coli* BTH101). In parallel, we constructed a genomic SPA library by partial digestion of SPA gDNA with Sau3AI and cloning gel-purified 0.8-5kb DNA fragments into pUT18. Transformation of the SPA library into *E. coli* BTH101 harboring pKT25::*yhjH*_SPA_ and plating on LB X-gal agar plates supplemented with Kanamycin (to select for pKT25::*yhjH*_SPA_) and ampicillin (to select for the SPA library cloned into pUT18) allowed us to screen ∼ 34,500 colonies that covered the SPA genomes about 7-fold. Overall, in this screen, we identified nine blue-stable colonies, harboring putative YhjH-interacting protein fusions. Sequencing the insert of these nine colonies (Table S1) indicated that three of them, which were constructed in two independent libraries, harbored a T18 fusion with the SPA gene *yciG* (accession number XBC74621).

**Figure 6.**
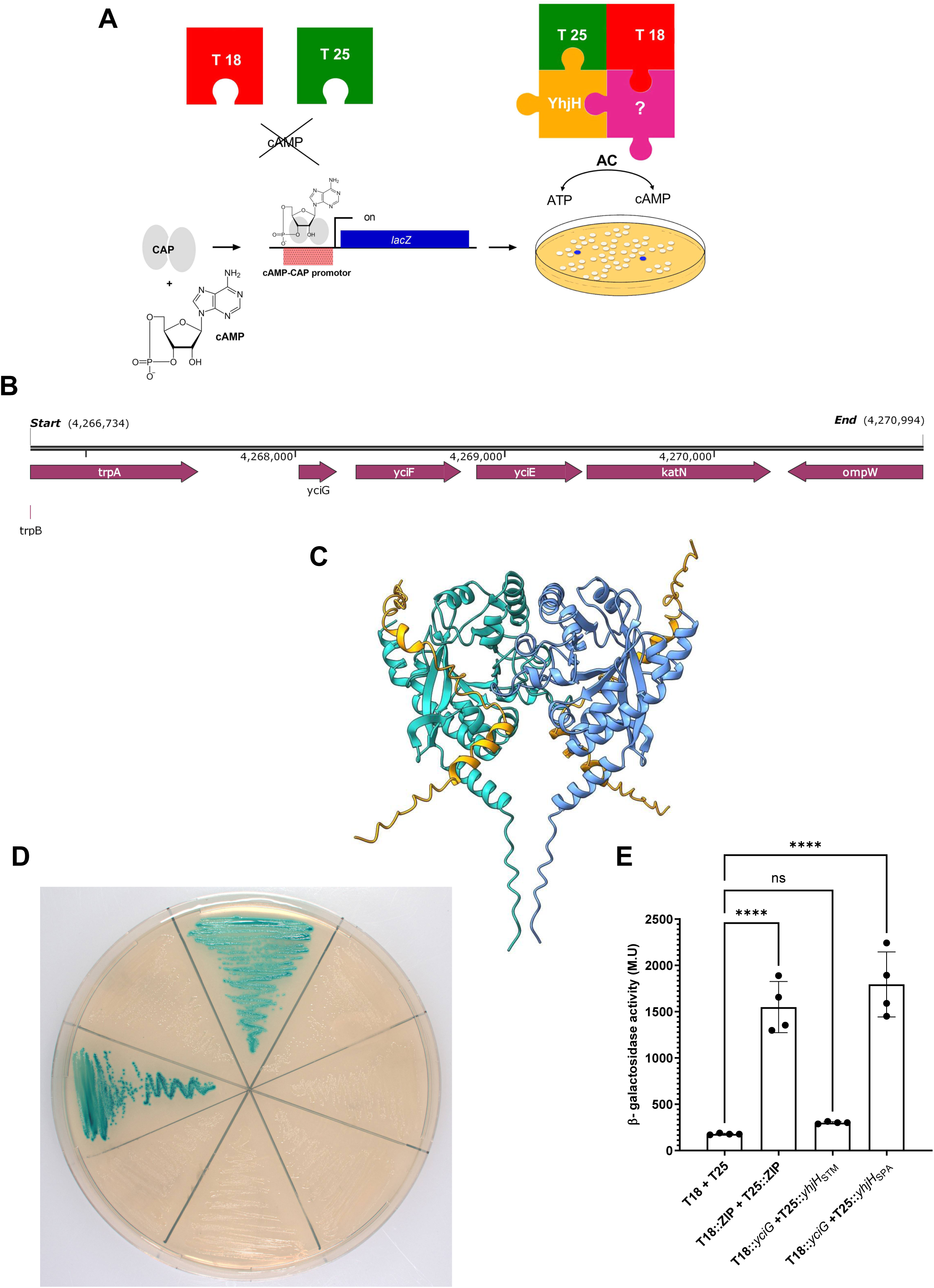
YhjH interacts with the general stress protein YciG in SPA but not in STM. (**A**) A schematic illustration of the BACTH-based genetic screen is shown. The catalytic domain of adenylate cyclase (CyaA) from *Bordetella pertussis* is separated into two sub-domains, T25 and T18, which are not active at their separated form. The sub-domain T25 was fused with YhjH_SPA_ and T18 was fused with a random genomic library of SPA. Heterodimerization of T25::*YhjH* with an interacting protein represented in the library results in a functional complementation of cAMP synthesis. When cAMP is being synthesized, it binds to the catabolite activator protein (CAP) and the cAMP/CAP complex induces the expression of lactose catabolic genes. These clones express a functional β-galactosidase enzyme (LacZ) and can be distinguished on LB agar plates in the presence of the chromogenic X-gal substrate as blue colored colonies. (**B**) The genetic organization of the *yciG* loci and its location as the first gene in the *yciGFE*-*KatN* operon is shown for the SPA 45157 genome. The genomic coordinates are shown at the beginning and end points. (C) An Alphafold model prediction showing YhjH dimer (in cyan and blue) and YciG (in gold) interaction. (**D**) *E. coli* BTH101 reporting strains containing paired combination of pUT18 and pKT25 plasmid derivatives were streaked on LB agar plate supplemented with X-gal and incubated at 30 °C for 48 h. BTH101 cells harboring pUT18C::ZIP and pKT25::ZIP expressing fused proteins to the leucine zipper domain of the yeast protein GCN4 were used as a positive control. Empty pUT18 and pKT25 plasmids harboring no insert were included as a negative control. (**E**) β-galactosidase activity assay was used to quantify YciG and YhjH interaction that is proportional to the activity of LacZ. *E. coli* BTH101 reporting strains containing the plasmid pairs pKT25::*yhjH*_SPA_ and pUT18C::*yciG*_SPA_, pKT25::*yhjH*_STM_ and pUT18C::*yciG*_SPA_, pUT18C::ZIP and pKT25::ZIP as a positive control, and empty pUT18 and pKT25 plasmids as a negative control were all grown at 30°C overnight followed by a colometric β-galactosidase assay. Bars show the mean and SEM of four independent repeats. Statistical significance was determined by one-way ANOVA followed by *post hoc* Dunnett’s test. ****, P-value < 0.0001; ns, not significant.

*yciG* is the first of the four-gene operon (*yciGEF-katN*; Fig. 6B), known to be regulated by RpoS and therefore is referred to as a general stress protein (44, 45). The gene *yciG* encodes a conserved 60 amino acid protein, which was previously shown to be involved in the regulation of flagellum-dependent motility in STM in an unknown manner (46).

To model possible protein–protein interaction between YhjH and YciG, the AlphaFold program (47) was used. This analysis indeed predicted that YhjH and YciG potentially interact in the c-di-GMP binding site of YhjH (Figure 6C).

To verify experimentally that YciG interacts with YhjH, the entire *yciG* ORF was cloned into pUT18 (creating the construct pUT18C::*yciG*) and introduced into BTH101 cells, expressing the STM or SPA allele of YhjH (pKT25::*yhjH*_STM_ or pKT25::*yhjH*_SPA_, respectively). Transformant BTH101 cells were plated on LB agar plates supplemented with X-gal under ampicillin and kanamycin selection and incubated at 30 °C. As a positive control, we used the two plasmids pUT18C::ZIP and pKT25::ZIP, in which the adenylate cyclase fragments, T25 and T18, are fused to the leucine zipper domain of the yeast protein GCN4 (42). These constructs are known to create a functional complementation mediated by the interaction of their leucine zipper motif (41), and resulted, as expected, in a robust β-galactosidase activity (Fig. 6D). In contrast, negative control strains expressing the constructs pUT18 and pKT25 alone or any other combination that expresses pKT25::*yhjH* or pUT18::*yciG* without their interacting partners did not generate any visible blue pigment under these conditions. Interestingly, using this BACTH system to assess the interaction of YhjH with YciG indicated that YciG interacts with YhjH_SPA_, but not with YhjH_STM_ (Fig. 6D).

Since protein–protein interaction in this system induces expression of the *lac* operon, and β-galactosidase activity is proportional to the degree of interaction, a quantitative β-galactosidase assay was performed to estimate the level of interaction between these proteins. In agreement with the qualitative results on LB X-Gal plates, β-galactosidase activity in strains expressing YhjH_SPA_ and YciG_SPA_ was approximately sixfold higher than that measured in strains expressing YhjH_STM_ and YciG_STM_ (Figure 6E). We concluded from these results that the small conserved protein YciG interacts with YhjH_SPA_, but its interaction with its STM ortholog is significantly weaker and therefore, YciG might play a different role in these serovars.

### The different interaction of YhjH with YciG in SPA vs. STM is not mediated by the YhjH sequence variation on the protein level

In contrast to YciG, which is fully conserved and possesses the same protein sequence in both serovars (Fig. 7A), YhjH varies in two amino acids between STM and SPA at positions 192 and 199, as shown earlier in Fig. 1A. Therefore, we speculated that the distinct interaction of YciG with the YhjH alleles in SPA vs. STM is driven by at least one of these two amino-acid variations. To test this hypothesis, site-directed mutagenesis was performed to replace positions P192 and T199 in YhjH_STM_ with A and I, respectively, yielding the same amino acid sequence as YhjH_SPA_. The modified *yhjH* alleles were cloned into the pKT25 plasmid and transformed into BTH101 cells harboring pUT18::*yciG*. Nonetheless, analyzing the potential interaction of the modified YhjH_STM_ mutants containing the substitutions P192A, T199I, or both with YciG, using the BACTH system, did not indicate that these changes increased YhjH_STM_ interaction with YciG, and that the T199I substitution may have even reduced YhjH interaction with YciG compared to the WT background (Figure 7B). We concluded from these results that the identified differences in the amino acid sequence of YhjH between SPA and STM are not responsible for the distinct interaction between YciG and YhjH in SPA vs. STM.

**Figure 7.**
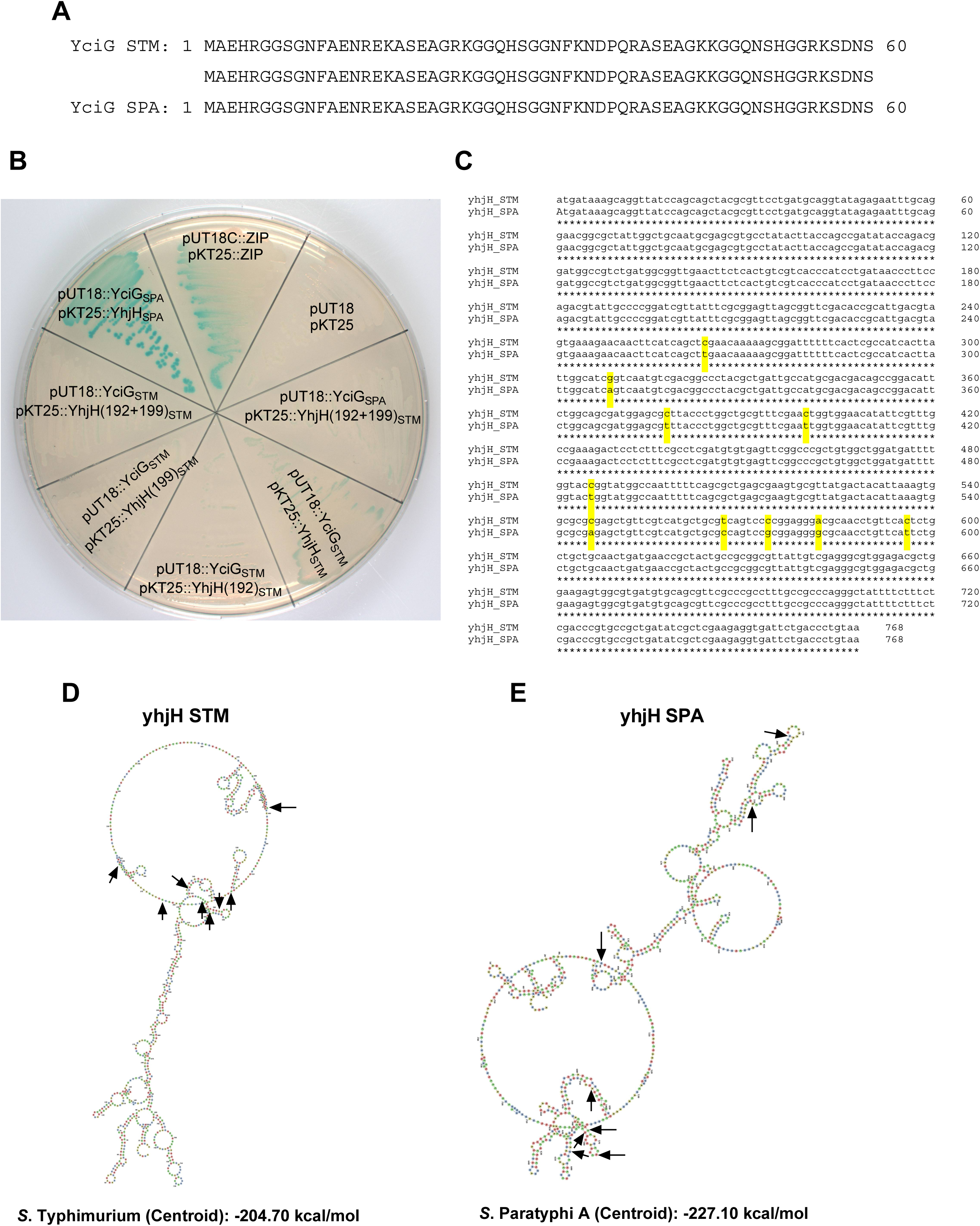
The different interaction of YhjH with YciG in SPA vs. STM is not mediated by sequence variation on the protein level. (**A**) A pairwise comparison of the YciG protein sequence in STM and SPA is shown to highlight its complete identity on the protein level. (**B**) A pairwise comparison of the *yhjH* gene sequence in STM and SPA is shown. Ten mismatches at the DNA level are highlighted in yellow. (**C**) *E. coli* BTH101 reporting strains containing paired combination of pUT18 and pKT25 plasmids with different inserts were streaked on LB agar plate supplemented with X-gal and incubated at 30 °C for 48 h. YhjH protein from STM with amino acid substitution at position 192 (P192A; YhjH(192)_STM_), 199 (T199I; YhjH(199)_STM_), or both (YhjH(192+199)_STM_) were included in this assay. BTH101 cells harboring pUT18C::ZIP and pKT25::ZIP were used as a positive control. Empty pUT18 and pKT25 plasmids were included as a negative control. (**D** and **E**) The predicted centroid secondary structure of the STM *yhjH* (D) and SPA *yhjH* (E) mRNA by RNAfold web server is shown. Nucleotide variations at the RNA level are indicated by arrows.

This has led us to consider an alternative mechanism, which is independent of the YhjH variation on the protein level and encouraged us to look more closely at sequence differences presented on the DNA or RNA level. Besides the non-synonymous mutations found in nucleotide 574 and 596, resulting in amino acid variation at position 192 and 199, respectively, eight additional synonymous variations were found in the *yhjH* gene sequence that maintain the same amino acids, but use different codons, with five of which being clustered in a high-density distribution around the amino acid changes (Fig. 7C and Table S2). To predict how these variations may affect the secondary RNA structure of *yhjH* we used the RNAfold server (48). Computational modeling of mRNA folding indicated that these synonymous changes dramatically affect RNA folding and despite 1.3% sequence difference at the DNA level, major mRNA reorganization was predicted with only 39% structural similarity between *yhjH* mRNA variants in SPA vs. STM (Fig. 7D-E). Moreover, centroid secondary structure predictions revealed a 22.4 kcal/mol stability difference (STM: -204.70 kcal/mol; SPA: -227.10 kcal/mol), indicating significantly enhanced mRNA stability of *yhjH* in SPA compared to its STM homolog. Based on these results, we raise the possibility that distinct mRNA folding may affect translation kinetics or a different protein conformation that alters interaction with YciG. Nonetheless, this prediction suggests that even with similar final amino acid sequences, the *yhjH* RNA could fold into different secondary structures that may affect its interaction with YciG in SPA vs. STM.

## DISCUSSION

Flagellar motility in *Salmonella* is an important virulence-associated function, controlled through a complex multi-layered regulatory hierarchy involving transcriptional, post-transcriptional, and post-translational mechanisms (49, 50). Although typhoidal and non-typhoidal salmonellae share many components of their regulatory program that orchestrates motility, we have previously identified curious differences in their motility regulation, expected to be relevant for their distinct lifestyle and pathogenesis (32–34). Our current work reveals a new machinery that shows how this sophisticated regulatory network has diverged between STM and SPA through gene inactivation and gain of function (neofunctionalization) of key regulatory components. While previous studies have documented the role of the PDE YhjH in STM motility and biofilm regulation (28, 29, 31, 35), our findings demonstrate that although the role of YhjH in biofilm formation is conserved, motility regulation by YhjH is entirely absent in SPA due to *ycgR* pseudogenization and the loss of c-di-GMP-mediated motility regulation in this serovar. In many Enterobacteriaceae members, c-di-GMP signaling provides a layer of post-translational control through the flagellar brake protein YcgR, which functions as a critical effector linking c-di-GMP levels to flagellar motor activity through direct protein-protein interactions with motor components upon c-di-GMP binding (25, 51). The truncation of YcgR into a nonfunctional peptide in SPA effectively severs this connection, rendering c-di-GMP level fluctuations inconsequential for motility control. This represents a fundamental rewiring of the conserved c-di-GMP regulatory circuit, with potential implications for how these pathogens respond to environmental signals and establish infection.

Genome degradation represents a common evolutionary trajectory among bacterial pathogens adapting to specialized niches, particularly those with restricted host ranges. This process has been extensively documented across diverse pathogenic lineages, including *Mycobacterium leprae*, *Rickettsia* species, *Yersinia pestis*, and typhoidal *Salmonella* serovars (52, 53). Previously, we and other groups have shown that in typhoidal *Salmonella*, approximately 5% of the genome consists of pseudogenes, compared to less than 1% in the broad-host-range STM (17, 18, 54). Here, we demonstrate how such genome degradation can fundamentally rewire regulatory networks, using the c-di-GMP signaling pathway as a paradigm for understanding the functional consequences of gene inactivation.

While previous studies have highlighted the loss of metabolic pathways and the downregulation of the motility regulon (55, 56), our current work identifies a specific "rewiring" of the c-di-GMP signaling network that alters the way SPA regulates its motility. The loss of YcgR by means of genome degradation effectively "liberates" the flagellar motor from one of its primary inhibitory signals, potentially allowing SPA to remain motile, even under conditions that would trigger biofilm formation or sessility in NTS serovars, in a way that may be critical for its unique intracellular lifestyle. Indeed, we recently demonstrated that a significant proportion of intracellular SPA cells are motile and reside within the host cell cytosol, unlike STM, which is largely vacuolar and non-motile (33). Lacking the YcgR brake likely facilitates this cytosolic motility, and its "always-ready" motility state may be essential for SPA to exit host cells and re-infect neighboring cells, thereby promoting systemic dissemination within the human host

The evolutionary forces driving genome degradation in host-restricted pathogens include reduced selective pressure for maintaining genes unnecessary in simplified environments, genetic drift in small effective population sizes, and potential direct selection for gene loss when certain functions become maladaptive (57). The results shown here suggest that rather than simply representing evolutionary decay, genome degradation can create novel regulatory architectures that may contribute to pathogen adaptation. This notion is further demonstrated by the serovar-specific interaction of YhjH with YciG that was found in SPA, but not in STM. While the primary role of YhjH is PDE activity, the identification of this interaction (that was hit three independent times in our genetic BACTH screen) and the presence of specific synonymous and non-synonymous mutations in the SPA *yhjH* sequence suggest that YhjH in SPA have gained a non-canonical role, which is mediated by its newly evolved interaction with YciG. Despite exhibiting more than 99% amino acid identity, YhjH demonstrates dramatically different regulatory roles in STM versus SPA. Both proteins retain similar c-di-GMP phosphodiesterase activity *in vitro*, yet only YhjH_STM_ regulates motility and only YhjH_SPA_ interacts with YciG.

The failure of site-directed mutagenesis targeting the two amino acid differences in YhjH to restore YciG interaction in STM was initially puzzling, but led us to consider alternative mechanisms. Our analysis revealed eight additional synonymous nucleotide changes in *yhjH* that maintain identical amino acids, but utilize different codons. Computational analysis using RNAfold predicted that these synonymous changes alter mRNA secondary structure and stability. These findings raised the intriguing possibility that synonymous mutations may lead to differences in mRNA structure that in turn influence protein function through effects on translation kinetics or co-translational folding processes. Such mechanisms could theoretically generate proteins with identical amino acid sequences, but subtly different cellular abundance or conformation capable of differential protein interactions. However, we emphasize that this represents a working hypothesis that still requires future direct experimental validation.

Nevertheless, the functional divergence of YhjH between serovars indicates that the regulatory differences arise from network-level changes and, potentially through serovar-specific interactions with YciG rather than protein-level alterations. The conservation of the PDE enzymatic activity alongside loss of motility regulatory function suggests that YhjH may serve alternative physiological roles in SPA. c-di-GMP signaling affects numerous cellular processes beyond motility, including biofilm formation, virulence gene expression, and cell cycle progression (21, 58). These diverse roles imply that c-di-GMP regulation remains important for other aspects of SPA physiology. The discovery that YhjH interacts with the general stress protein, YciG in SPA represents a potentially significant finding that warrants further investigation. YciG is part of the RpoS-regulated stress response (45) and has previously been shown to be involved in the transcriptional regulation of *fliC* (46). A null *yciG* STM mutant was also reported to present attenuated survival under acidic conditions and inside macrophages, impaired adhesion and invasion into epithelial cells, and reduced swimming motility (59). The interaction between a PDE and a stress-response protein suggests a regulatory link between c-di-GMP signaling and the general stress response, which may be particularly relevant during the transition from the gut environment to the intracellular niche. However, the physiological significance of this interaction and its role in SPA pathogenesis remain to be determined through future functional studies.

Overall, our findings illustrate how genome degradation in bacterial pathogens can extend beyond simple gene loss to encompass complex regulatory network reorganization. The loss of YcgR-mediated motility control, coupled with the potential gain of YciG interaction, suggests that SPA has evolved alternative regulatory mechanisms that may be better suited to its lifestyle as a human-restricted systemic pathogen. The differential motility regulation patterns observed may reflect the distinct ecological niches occupied by STM and SPA. STM’s robust environmental motility responses may be optimized for survival in diverse environments and rapid colonization of multiple host species. In contrast, SPA’s simplified motility regulation, combined with its demonstrated intracellular motility (33), may be better adapted for systemic infection and intracellular survival in human hosts (Fig. 8).

**Figure 8.**
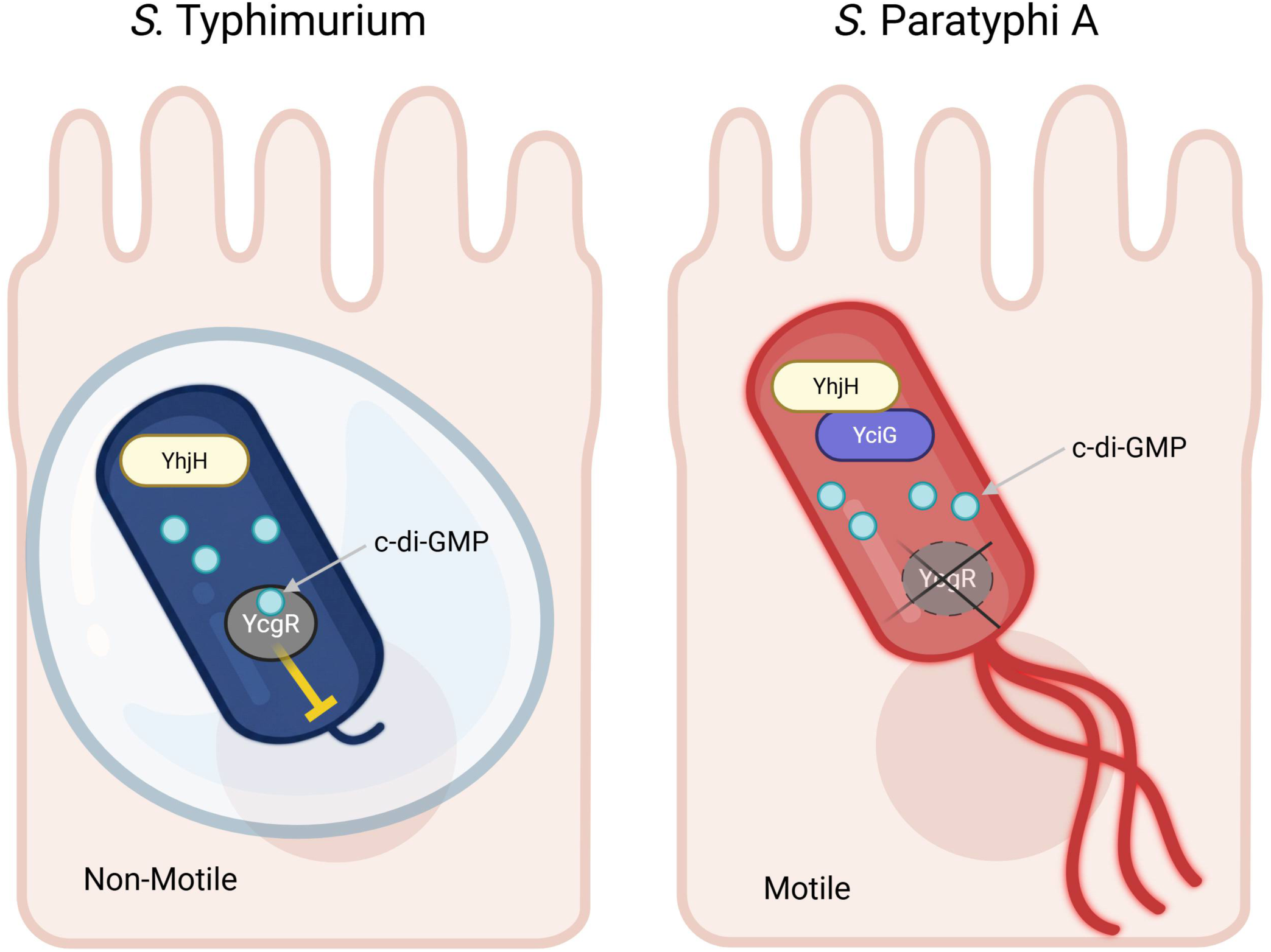
A Working model for the different role of YhjH in STM vs. SPA. In both STM and SPA serovars YhjH functions as PDE to degrade c-di-GMP and keep its intracellular level low. In STM, c-di-GMP binds to the flagellar brake protein YcgR to slow down the flagella motor speed in response to high c-di-GMP levels, resulting in suppression of the motility. In SPA, YcgR is not expressed due to pseudogenization and therefor c-di-GMP signaling does not control flagella-mediated motility in a manner that may maintain SPA locomotion under normally sessility-favoriting conditions, such as intracellular infection. Instead, serovar-specific protein interaction between YhjH and the general stress protein YhjH has evolved in SPA, but not in STM, implying a new role of YhjH in stress tolerance.

In summary, our work provides a molecular basis for the altered motility regulation in two clinically important *S. enterica* serovars. The degradation of *ycgR* in SPA represent a significant pathoadaptive shift. By dismantling the YcgR-mediated brake and ensuring a low c-di-GMP environment, SPA has optimized its flagellar function for a systemic, intracellular lifestyle. These findings not only highlight regulatory differences in the motility of typhoidal and non-typhoidal *Salmonella*, but also underscore the importance of c-di-GMP signaling as a target for "genome degradation" during the evolution of a human-restricted pathogen. Comparative analysis of c-di-GMP signaling systems across other host-adapted bacterial pathogens would help determine how widespread such regulatory network degradation may be and whether common evolutionary patterns exist.

## MATERIALS AND METHODS

### Bacterial strains and growth conditions

All bacterial strains and plasmids used in this study are listed in Table S3. *S*. Typhimurium SL1344 (60) and *S*. Paratyphi A 45157 (61) were used as the parental strains, and all of the constructed mutants were isogenic strains derived from these backgrounds. *Salmonella* and *E. coli* strains were cultured in LB (Lennox) medium at 30 or 37 °C. Liquid cultures were grown on a roller drum or in Erlenmeyer flasks incubated in a shaker incubator with a shaking speed of 200 RPM. When appropriate, the medium was supplemented with antibiotics in the following concentrations: kanamycin, 50 µg/mL; ampicillin, 100 µg/mL; chloramphenicol 10 μg/mL; anhydrotetracycline (AHT) 500 ng/mL.

### Site directed mutagenesis and construction of null deletion strains

All primers used in this study are listed in Table S4. Oligonucleotides were purchased from IDT and PCR was carried out using Phusion Hot Start Flex DNA Polymerase (New England BioLabs) for cloning or with Red Load Taq Master (Larova) for confirmatory PCR. None-polar null deletions of *yhjH* (Δ*yhjH*)*, ycgR* (Δ*ycgR*), and the double mutant strain Δ*yhjH* Δ*ycgR* were constructed using the λ Red–mediated recombination system, while the resistance cassette was subsequently eliminated from the genome by using a helper plasmid encoding the FLP recombinase as previously described (62).

Site directed mutagenesis was conducted as described in (63). For replacing the glutamate at position 136 to alanine (E136A) in the *yhjH* gene in STM, the primers ‘yhjH E136A TC1 F’ and ‘yhjH E136A TC1 R’ were first used to amplify an I-SceI cleavage site together with a kanamycin resistance cassette from the template vector pWRG717, producing the first targeting construct (TC1) for recombination. The obtained amplicon was transformed into heat-shock induced electrocompetent STM cells containing pWRG730 and recombinant transformants were selected on LB plates containing 10 μg/mL chloramphenicol and 25 μg/mL kanamycin following O/N incubation at 30 °C. For the second recombination step, the second targeting construct (TC2) was generated by PCR using STM gDNA as a template and the primers ‘yhjH E136A TC2 F’ and ‘yhjH E136A TC2 R’. The TC2 amplicon was then transformed into heat-shock induced electrocompetent STM harboring pWRG730 and an integrated kanamycin resistance cassette in the target site. Electroporated cells were then plated on LB agar plates containing 10 μg/mL chloramphenicol and 500 ng/mL anhydrotetracycline, and were incubated O/N at 30 °C. To remove the recombinase (pWRG730 curing) Cm^r^ Kan^s^ PCR-verified colonies were streaked on LB plates and grown at 42° C for O/N.

To create an E136A YhjH substitution in SPA the primers ‘yhjH E136A TC1 F SPA’ and ‘yhjH E136A TC1 R’ were used to amplify the TC1 construct from the template vector pWRG717 and the primers ‘yhjH E136A TC2 F SPA’ and ‘yhjH E136A TC2 R SPA’ were used to amplify the TC2 construct using SPA gDNA as above. Final substitutions in the genome were verified by Sanger sequencing of a PCR amplicon obtained using primers flanking the *yhjH* gene (‘seq yhjH F STM/SPA’ and ‘seq yhjH R STM/SPA’).

Scarless point mutations (C574G and C596T) were introduced into the *yhjH* gene of STM, resulting in the amino acid substitutions P192A and T199I, respectively. The resistance cassette derived from pWRG717 was amplified using the primers ‘TC1 *yhjH* 192 F’ and ‘TC1 yhjH 192 199 R’ for the P192A (C574G) substitution, and the primers ‘TC1 *yhjH* 199 F’ and ‘TC1 yhjH 192 199 R’ for the T199I (C596T) substitution. To amplify the TC2 constructs, the primers ‘TC2 yhjH 192 F’ and ‘TC2 yhjH 192 199 R’ were used for the P192A substitution, and the primers ‘TC2 yhjH 199 F’ together with ‘TC2 yhjH 192 199 R’ for the T199I substitution were used. In both cases, the STM SL1344 gDNA was added to the PCR reaction as a template. Selection of the second recombination event (to eliminate the resistance cassette) and pWRG730 curing was done as above.

### Genetic complementation and molecular cloning

For complementation of the Δ*yhjH* deletion in SPA and STM, the corresponding sequences were PCR amplified using gDNA from WT *S*. Paratyphi A 45157 and *S*. Typhimurium SL1344 ,respectively, as a template and the primers ‘Clone yhjH ClaI F’ and ‘Clone yhjH XmaI R’. The obtained amplicons that included the *yhjH* ORF and its upstream regulatory region were digested with ClaI and XmaI and cloned into the low-copy number plasmid pWSK29.

*yjcC* cloning was conducted by PCR amplification using *S*. Typhimurium SL1344 gDNA as a template and the primers ‘yjcC xbaI-F’ and ‘yjcC hind-R’. The obtained amplicon was digested with XbaI and HindIII and cloned into pBAD18 under arabinose induced promoter.

Complementation of the Δ*ycgR* deletion was achieved by PCR amplification of the entire *ycgR* ORF, including its upstream regulatory sequence, using *S*. Paratyphi A 45157 and *S*. Typhimurium SL1344 genomic DNA as templates with the primers ‘ycgR region-sacI-F’ and ‘ycgR-bamHI-R’. The obtained amplicons were digested with ScaI and BamHI and cloned into pWSK29.

To study YcgR expression on the protein level, its corresponding coding sequence from STM and SPA was PCR amplified using the primers ‘pet28AycgR-F’ and ‘pet28AycgR-R’ and cloned using the Gibson Assembly into the pET28a plasmid that was transformed into electrocompetent *E. coli* BL21 cells. The obtained constructs express an N-terminus 6×His -tagged version of YcgR from STM and SPA, under an IPTG-inducible promoter.

To study the protein-protein interaction of YhjH and YciG using the BACTH system (see below) the intact sequences of *yhjH* and *yciG* were PCR amplified using SPA or STM gDNA as a template. *yhjH* was amplified using the primers ‘*yhjH* BamHI F’ and ‘*yhjH* SmaI R’, while *yciG* was amplified using the primers ‘*yciG* BamHI F’ and ‘*yciG* SmaI R’. The obtained PCR products resulted in 765 and 177 bp amplicons, respectively were digested with BamHI and SmaI and cloned into pKT25 and pUT18, respectively.

Similarly, the modified P192A and T199I *yhjH* alleles were amplified by PCR from STM gDNA using the primers “*yhjH* BamHI F” and “*yhjH* SmaI R”, digested with BamHI and SmaI, respectively, and cloned into the pKT25 plasmid.

### Expression and purification of His-tagged proteins

The *yhjH* gene from STM and SPA was amplified using the primers ‘yhjH his-F-NdeI’ and ‘yhjH his-R-BamHI’ and *Salmonella* gDNA as a template. The obtained amplicons were digested with NdeI and BamHI and cloned into the pET28a vector, generating a C-terminus 6×His tagged YhjH, under an IPTG inducible promoter. The resulted constructs (pET28a::*yhjH*_STM_ and pET28a::*yhjH*_SPA_) were transferred into electrocompetent *E. coli* BL21 and selected on LB agar plates supplemented with kanamycin. To produce the YhjH-His tagged proteins, overnight cultures of *E. coli* BL-21 harboring the constructs were grown under kanamycin selection at 37 °C with shaking, subcultured 1:100 into 3 mL of fresh LB and grown for 2 h to an OD_600_ of ∼0.6. IPTG was added to the cultures to a final concentration of 1 mM, and the induced cultures were continued to grow for an additional 4 h under the same conditions. To harvest the cells, the cultures were placed on ice for 10 min and centrifuged for 5 min at 5,000 *g* at 4 °C. The bacterial pellets were resuspended in 500 µL of Lysis Buffer (50 mM NaH _2_PO _4_, 300 mM NaCl, 10 mM imidazole, pH 8.0) supplemented with 50 µL of 10 mg/mL lysozyme and 9 U of Benzonaze endonuclease and incubated for 30 min at 4 ^°^C with gentle rotation.

To release the YhjH-His from the insoluble fraction, 50 µL of 10 % n-dodecyl-β-D-maltoside (DDM; Sigma-Aldrich) detergent was added, and the lysate was further incubated for an additional 90 min under the same conditions. After 2 h, 500 µL of Binding/Washing Buffer of the MagListo His-tagged protein purification kit (Bioneer) were added to the lysate, and purification on nickel-beads was carried out according to the manufacturer’s instructions, while both binding and elution steps were extended to 1 h at 4 °C under gentle shaking. The purified proteins, together with analytical aliquots from each purification step, were separated on 12% SDS-PAGE and analyzed by Coomassie Brilliant Blue R250 staining. Protein concentrations were determined using the Pierce 660nm Protein Assay (Thermo Fisher Scientific) compatible with imidazole, according to manufacturer’s manual.

### Analysis of c-di-GMP metabolites

WT and Δ*yhjH* isogenic strains of STM and SPA were subcultured (1:50) into fresh LB medium (10 mL) and grown for 8 h at 30 °C under aerobic conditions. Bacterial cultures were centrifuged at 2,500 *g* for 20 min at 4 °C, and washed twice with 500 µL of fresh LB. The washed pellets were resuspended with 300 μL of ice-cold extraction solvent of acetonitrile/methanol/water (2/2/1, v/v/v) and incubated on ice for 15 min. Bacterial extracts were subsequently incubated for 10 min at 95 °C to inactivate phosphodiesterases and cooled down on ice. The bacterial extracts were then centrifuged at 20,800 *g* for 10 min at 4 °C, and the supernatants were extracted twice again with 200 μL of extraction solvent. The extracted samples were centrifuged at 20,800 *g* for 10 min at 4 °C, and the supernatants were transferred into new 2.0 mL tubes, which were evaporated to dryness using a Speed-Vac vacuum concentrator at 40 °C. Bacterial extracts were kept at -80 °C until analysis at the Research Core Unit Metabolomics of the Hannover Medical School. For c-di-GMP detection and quantitation, samples were resolved in 150 μL ddH_2_O and analyzed by High-Performance Liquid Chromatography-Tandem Mass Spectrometry (HPLC-MS/MS) on a QTrap5500 triple quadrupole mass spectrometer (Sciex, Framingham, MA, United States) as described before (64).

### Measuring enzymatic activity of phosphodiesterase

c-di-GMP phosphodiesterase activity was estimated by monitoring the change in the fluorescence of a cyclic di-GMP subtract (2’-, 2’’- O- (Di-N’-methylanthraniloyl)-cyclic diguanosine monophosphate) that was MANT-labeled (Di-MANT-c-di-GMP), as described in (65) with slight modifications. Nickel-purified YhjH-His was concentrated 2-fold by a Speed-Vac vacuum concentrator at room temperature, resuspended in 0.4 volumes of Preservation Buffer (70% Glycerol; 70 mM TRIS pH 8.0, 80 mM KCl, 90 mM MgCl_2_, 4 mM DTT), and kept at -80 °C. PDE activity was measured using 1.2 µg (∼ 42 pmol) protein in a 100 µL reaction solution (50 mM Tris-HCl, 250 mM NaCl, 25 mM KCl, 5 mM MgCl_2_, 1 mM DTT, 0.5 μM MANT–c-di-GMP). The decrease in fluorescence at 440 nm was measured in a white 96-well flat-bottom plate (Greiner) at 37 °C using the Infinite 200 Pro M-PLEX microplate reader (Tecan, Switzerland). After the background fluorescence was subtracted from the wells without MANT–c-di-GMP, the remaining fluorescence over time was calculated as a percentage of the fluorescence at time point zero. MANT-c-di-GMP only (without protein) was included as a control for spontaneous degradation.

### Motility assay

Swimming motility was determined as described before (56). Briefly, *Salmonella* cultures were grown overnight at 37 °C, diluted 1:100 in fresh LB medium, and grown for an additional three hours under the same conditions. Aliquots of 10 µL were placed in the center of soft LB agar plates and incubated for 3-5 hours at 37 °C without being inverted. The motility radius was determined with a measuring ruler and imaged using a Fusion Solo X system (Vilber).

### Biofilm formation

*Salmonella* SPA and STM cultures were grown in LB-Lennox broth for 16 h at 37 °C and subcultured 1:100 into fresh LB medium lacking sodium chloride (10 g/L peptone, 5 g/L yeast extract). Aliquots of 150 µl from each culture were placed into 96-well cell-culture-treated microplates and incubated at 28 °C without shaking. After 96 hours, planktonic cells were removed and the attached biofilm layer was fixed for 2 h at 60 °C, following by staining with 150 μl of 0.1% Crystal Violet dye for 10 min at room temperature. After washing with Phosphate-Buffered Saline (PBS), the stained biofilm was resuspended in 150 µl of acetic acid (33%) and biofilm quantification was achieved by measuring the optical density at 560 nm of the crystal violet-stained biomass using the Infinite M Plex (Tecan) plate reader.

### Identification of YhjH Interacting proteins by a BACTH-based genetic screening

Genetic screening for YhjH interacting proteins was performed using a bacterial adenylate cyclase two-hybrid system (BACTH) approach (43), adapted from the Euromedex BACTH system kit’s protocol (https://shopresearch.euromedex.com/document/FTPDF/EUK001.pdf).

Genomic DNA (gDNA) from *Salmonella* Paratyphi A 45157 was extracted using the GenElute Bacterial Genomic DNA Kit (Sigma-Aldrich). The gDNA was partially digested with Sau3AI (Thermo Fisher Scientific) and separated by electrophoresis on a 0.8% agarose gel. To construct dense libraries, three independent fragment size ranges (0.8-1.5; 1.5-3; and 3-5 kb) were excised and gel-purified using the QIAEX II Gel Extraction Kit (QIAGEN). For library cloning, the BACTH prey vector, pUT18 was digested with BamHI (New England Biolabs) at 37 °C for 2 h and dephosphorylated with Shrimp Alkaline Phosphatase (Thermo Fisher Scientific) at 37 °C for 1 h. Each of the three independent gel-purified DNA fragment pools was ligated overnight at 4 °C into the linearized dephosphorylated pUT18 vector using T4 DNA ligase (Promega) at an insert: vector molar ratio of 6:1.

The resulting ligation products were transformed into electrocompetent *E. coli* MC1022 cells by electroporation. Transformed cells were plated on LB-agar plates containing ampicillin (100 µg/mL) and incubated at 37 °C overnight. The obtained transformants from each library were collected by scraping the grown colonies from the plates, pooled together, and subjected to plasmid isolation using the AccuPrep Plasmid Extraction Kit (Bioneer) to generate three independent pUT18-borne SPA genomic libraries.

To generate the T25-YhjH bait construct, the coding sequence for YhjH from SPA was cloned into the pKT25 vector, as detailed in the ’Genetic complementation and cloning’ section. For the screening, 10-20 ng of plasmid DNA from each pUT18 SPA genomic library was independently transformed into 60 μL of electrocompetent *E. coli* BTH101 reporter cells already carrying the pKT25::YhjH bait plasmid. Transformed cells were plated on large (150 mm) LB-agar plates supplemented with X-gal (40 µg/mL), IPTG (0.1 mM), ampicillin (100 µg/mL), and kanamycin (50 µg/mL) and incubated at 30 °C for 72 h.

Under these conditions, bacteria expressing interacting YhjH-prey hybrid proteins formed blue colonies due to adenylate cyclase activity, while cells expressing non-interacting proteins remained white. Putative positive interacting clones (blue colonies) were individually picked, and their pUT18 plasmids were isolated using AccuPrep Plasmid Extraction Kit (Bioneer). The genomic inserts within these plasmids were then sequenced using Sanger sequencing with the pUT18-specific primers ‘pUT18 check F’ and ‘pUT18 check R’ to identify the SPA genes encoding the YhjH-interacting protein fragments.

### β-galactosidase assay

*E. coli* BTH101 reporter strains harboring the different pUT18 and pKT25 constructs were grown overnight in LB supplemented with ampicillin and kanamycin at 30 °C. To determine the β galactosidase specific activity 500 µl of cultures were added to 500 µl Z-Buffer (0.06 M Na_2_HPO_4_·7H_2_O, 0.04 M NaH_2_PO_4_·H_2_O, 0.01 M KCl, 0.001 M MgSO_4_·7H_2_O, and 0.05 M β-mercaptoethanol; pH 7) and assayed as detailed in (66), using o-nitrophenyl-β-D-galactopyranoside (ONPG, Sigma-Aldrich) as a substrate.

### Sequence alignment, RNA folding modeling and protein’s 3D structure prediction

DNA and protein sequences alignment were conducted using the Clustal Omega Multiple Sequence Alignment tool at the EMBL-EBI portal (https://www.ebi.ac.uk/jdispatcher/msa/clustalo) (67). Prediction of mRNA folding was conducted using the RNAfold web server (http://rna.tbi.univie.ac.at) and prediction of proteins 3D structure and interactions was done using the AlphaFold Server (https://alphafoldserver.com/welcome).

## ACKNOWLEDGEMENTS

We are thankful to Dr. Lori Burrows for her generous gift of bacterial strains used for the BACTH experiment. The work at the Gal-Mor laboratory was supported by the Science Foundation (ISF) under grant number 1228/23 awarded to O.G.M. The funder has no role in study design, data collection and analysis, decision to publish, or preparation of the manuscript.

